# SECEDO: SNV-based subclone detection using ultra-low coverage single-cell DNA sequencing

**DOI:** 10.1101/2021.11.08.467510

**Authors:** Hana Rozhoňová, Daniel Danciu, Stefan Stark, Gunnar Rätsch, André Kahles, Kjong-Van Lehmann

## Abstract

**Motivation:** Several recently developed single-cell DNA sequencing technologies enable whole-genome sequencing of thousands of cells. However, the ultra-low coverage of the sequenced data (*<* 0.05x per cell) mostly limits their usage to the identification of copy number alterations in multi-megabase segments. Many tumors are not copy number-driven, and thus single-nucleotide variant (SNV)-based subclone detection may contribute to a more comprehensive view on intra-tumor heterogeneity. Due to the low coverage of the data, the identification of SNVs is only possible when superimposing the sequenced genomes of hundreds of genetically similar cells. Thus, we have developed a new approach to efficiently cluster tumor cells based on a Bayesian filtering approach of relevant loci and exploiting read overlap and phasing.

**Results:** We developed Single Cell Data Tumor Clusterer (SECEDO, lat. ‘to separate’), a new method to cluster tumor cells based solely on SNVs, inferred on ultra-low coverage single-cell DNA sequencing data. We applied SECEDO to a synthetic dataset simulating 7,250 cells and eight tumor subclones from a single patient, and were able to accurately reconstruct the clonal composition, detecting 92.11% of the somatic SNVs, with the smallest clusters representing only 6.9% of the total population. When applied to four real single-cell sequencing datasets from a breast cancer patient, each consisting of ≈2,000 cells, SECEDO was able to recover the major clonal composition in each dataset at the original coverage of 0.03x, achieving an ARI score of ≈0.6. The current state-of-the-art SNV-based clustering method achieved an ARI score of ≈0, even after increasing the coverage *in silico* by a factor of 10, and was only able to match SECEDO’s performance when pooling data from all four datasets, in addition to artificially increasing the sequencing coverage by a factor of 7. Variant calling on the resulting clusters recovered more than twice as many SNVs as would have been detected if calling on all cells together. Further, the allelic ratio of the called SNVs on each subcluster was more than double relative to the allelic ratio of the SNVs called without clustering, thus demonstrating that calling variants on subclones, in addition to both increasing sensitivity of SNV detection and attaching SNVs to subclones, significantly increases the confidence of the called variants.

**Availability:** SECEDO is implemented in C++ and is publicly available at https://github.com/ratschlab/secedo.

## 1 Introduction

Somatic single-nucleotide variants (SNVs) are commonly associated with cancer progression and growth (Stratton *et al*., 2009). The recent development of single-cell DNA sequencing technologies (Gawad *et al*., 2016) offers the ability to study somatic SNVs at a single-cell level, providing much more detailed information about tumor composition and phylogeny than traditional bulk sequencing (Kuipers *et al*., 2017; Navin *et al*., 2011). However, several technical obstacles decrease the interpretability of the data obtained using these technologies. In particular, most of the current single-cell DNA sequencing technologies require a whole-genome amplification step, which introduces artifacts such as DNA-amplification errors, allelic drop-out, imbalanced amplification, etc. (Gawad *et al*., 2016). Several approaches (Bohrson *et al*., 2019; Dong *et al*., 2017; Hård *et al*., 2019; Lähnemann *et al*., 2021; Luquette *et al*., 2019; Singer *et al*., 2018; Zafar *et al*., 2016) have been proposed to detect SNVs based on such data.

Approaches that do not require whole-genome amplification have been developed to overcome issues related to amplification (Laks *et al*., 2019; Navin *et al*., 2011). A prominent example of such technologies is 10X Genomics’ Chromium Single Cell CNV Solution^1^. This technology allows the sequencing of hundreds to thousands of cells in parallel, albeit with only extremely low sequencing coverage (*<*0.05x per cell). Hence, its use has been limited to the inference of copy number variations (CNVs) and alterations (CNAs) (10X Genomics, 2018; Durante *et al*., 2020; Velazquez-Villarreal *et al*., 2020; Zaccaria and Raphael, 2021). The attempts to also use these data for the identification of tumor subclones based solely on SNVs have so far failed to provide a solution that would be able to recover the clonal composition at the original sequencing depth (Myers *et al*., 2020); in particular, SBMClone, the algorithm of Myers *et al*. (2020), requires a minimum coverage of ≥ 0.2x per cell, roughly four times more than what is currently achievable using the 10X Genomics technology (10X Genomics, 2018; Velazquez-Villarreal *et al*., 2020).

In this work, we propose SECEDO, a novel algorithm for clustering cells based on SNVs using single-cell sequencing data with ultra-low coverage. Using an extensive set of simulated data, as well as four real data sets, we show that SECEDO is able to correctly identify tumor subclones in data sets with per-cell coverage as low as 0.03x, improving the current state of the art by a factor of seven and thus rendering the algorithm applicable to currently available single-cell data. We also provide an efficient C++ implementation of SECEDO, which is able to quickly cluster sequencing data from thousands of cells while running on commodity machines.

## 2 Methods

### Overview

Due to the extremely low coverage of the data (< 0.05x per cell), deciding whether two cells have identical or distinct genotypes is a difficult problem. Most loci are covered, if at all, by only one read (**Supplementary Figure S1**). This makes it difficult, if not impossible, to interpret an observed mismatch when comparing data from two cells. The mismatch could be caused by an actual somatic SNV, by a sequencing error, or by a heterozygous locus that was sequenced in different phase in the two cells. Hence, it is crucial to jointly leverage the information from all cells at the same time.

The pivotal blocks in the SECEDO pipeline (**Figure 1**) are (1) a Bayesian filtering strategy for efficient identification of relevant loci and (2) derivation of a global cell-to-cell similarity matrix utilizing both the structure of reads and the haplotype phasing, which proves to be more informative than considering only one locus at a time.

**Fig. 1.**
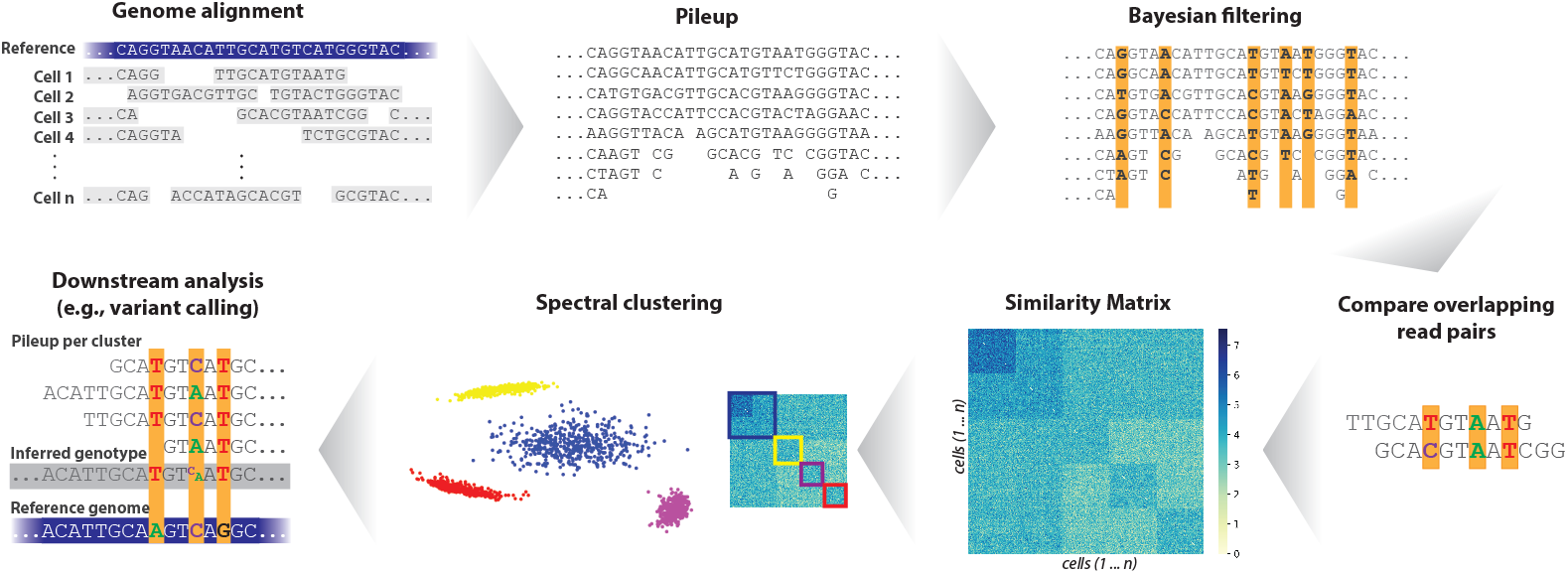
The SECEDO pipeline. After sequencing, reads are piled up per locus and a Bayesian filter eliminates loci that are unlikely to carry a somatic SNV. For each pair of reads, SECEDO compares the filtered loci and updates the likelihoods of having the same genotype and of having different genotypes for the corresponding cells. The similarity matrix, computed as described in Methods, is then used to cluster the cells into 2 to 4 groups (the number of groups depends on the data and is determined automatically by SECEDO) using spectral clustering. The algorithm is then recursively applied to each cluster until a termination criterion is reached.

SECEDO first performs a filtering step, in which it examines the pooled sequenced data for each locus and uses a Bayesian strategy to eliminate loci that are unlikely to carry a somatic SNV. The filtering step drastically increases the signal-to-noise ratio by reducing the number of loci by 3 to 4 orders of magnitude (depending on the coverage), while only eliminating approximately half of the loci that carry a somatic SNV. Moreover, the eliminated mutated loci typically have low coverage or high error rate and would not be very useful for clustering. In the second step, SECEDO builds a cell-to-cell similarity matrix based only on read-pairs containing the filtered loci, using a probabilistic model that takes into account the probability of sequencing errors, the frequency of SNVs, the filtering performance, and, crucially, the structure of the reads, i.e. the fact that the whole read was sampled from the same haplotype. In the third step of the pipeline, we use spectral clustering to divide the cells into two or more groups. At this point, we reduced the problem to an instance of the well-studied community detection problem (Porter *et al*., 2009), so spectral clustering is a natural choice. Optionally, the results of spectral clustering can be further refined in a fourth step using the Expectation-maximization algorithm (Dempster *et al*., 1977). The whole pipeline is then repeated for each of the resulting subclusters. The process is stopped if (1) there is no evidence for the presence of at least two clusters in the similarity matrix, or (2) the clusters are deemed too small. Downstream analysis, for instance, variant calling, can then be performed by pooling sequencing data from all cells in one cluster based on the results of SECEDO to create a pseudo-bulk sample.

### Filtering uninformative loci

Consideration of all genomic loci is not desirable when performing the clustering and variant calling, since most positions are not informative for clonal deconvolution. The most informative loci with respect to the clustering of the cells are the loci carrying somatic SNVs since they provide (1) information on assignment of cells to clusters and (2) information on haplotype phasing (due to loss/gain of heterozygosity). To a lesser extent, this is also true for germline heterozygous loci since they provide information on haplotype phasing. In other words, loci at which all the cells have the same homozygous genotype do not provide any information relevant to the task of dividing the cells into genetically homogeneous groups, so they can be excluded from downstream analysis.

Due to the low sequencing coverage, it is generally not possible to reliably assign genotypes to individual cells. However, we identify loci of interest by using the *pooled data* across all the cells to approximate posterior probabilities that the cells have the same genotype. Consider for example a specific locus at which all cells have genotype AA. Assuming sequencing errors happen independently with probability *θ* and are unbiased (i.e. all types of substitutions are equally probable), the fraction of As in the pooled data is in expectation (1 − *θ*) and the fraction of all other bases is *θ*/3. A locus with a significantly different proportion of observed bases indicates that there may be two (or more) different genotypes contributing to the observed data. In particular, we compute the posterior probability that all cells at the locus share the same homozygous genotype using an approximate Bayesian procedure. If this posterior is lower than a chosen threshold *K*, the locus is marked as ‘informative’.

Formally, let *C*_1_, *C*_2_, *C*_3_, *C*_4_ be the bases sorted from the most to the least frequent in the pooled data at the given position, *c*_1_, *c*_2_, *c*_3_, *c*_4_ the corresponding counts (*c*_1_ ≥ *c*_2_ ≥ *c*_3_ ≥ *c*_4_), *c* the total coverage (*c* = *c*_1_ + *c*_2_ + *c*_3_ + *c*_4_). Next, let *M* be an indicator random variable that is 1 if all cells in the sample have the same homozygous genotype and 0 otherwise. Applying Bayes rule, we can compute *P* (*M* | *c*_1_, *c*_2_, *c*_3_, *c*_4_) as:

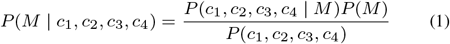

We compute or approximate the individual terms as follows:

- *P* (*M*) can be estimated from literature: the prevalence of somatic SNVs in cancer lies between 10^−9^ and 10^−3^ (Alexandrov *et al*., 2013; Lawrence *et al*., 2013); the frequency of heterozygous sites in a typical human genome lies between ca 0.04 and 0.11% (Bryc *et al*., 2013; Meyer *et al*., 2012). In order to be conservative, we choose the largest probability (≈ 10^−3^) in both cases, resulting in *P* (*M*) ≈ 1 − 2· 10^−3^ = 0.998.
- *P* (*c*_1_, *c*_2_, *c*_3_, *c*_4_ | *M*), is equal to

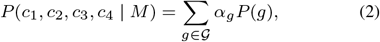

where *α*_*g*_ = *P* (*c*_1_, *c*_2_, *c*_3_, *c*_4_ | genotype of all cells is *g*) and 𝒢 = {*AA, CC, GG, TT*} is the set of all possible homozygous genotypes. The probability *α*_*g*_ of observing data (*c*_1_, *c*_2_, *c*_3_, *c*_4_) given that the genotype of all cells is *g* (*g* = *C*_*i*_*C*_*i*_) has a multinomial distribution with *c* trials and event probabilities equal to 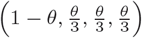:

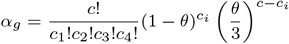

Assuming the error rate *θ* is small, the result of the equation above is negligible for any *c*_*i*_ that is not close to *c*. As a consequence, if the prior *P* (*g*) is approximately the same for all genotypes, we can approximate the sum in **Equation 2** with the largest term:

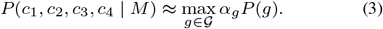

- Computing *P* (*c*_1_, *c*_2_, *c*_3_, *c*_4_) is intractable, as it would involve summing over all possible combinations of the cells’ genotypes. We instead approximate the evidence by

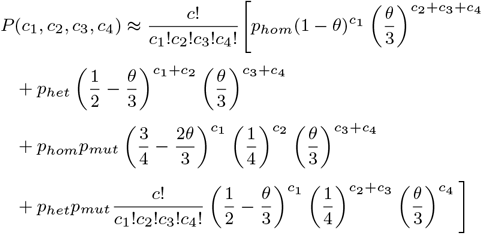

where *p*_*hom*_, *p*_*het*_, *p*_*mut*_ represent the probability of a locus being homozygous, heterozygous and mutated, respectively. The first summation term estimates *P* (*c*_1_, *c*_2_, *c*_3_, *c*_4_) for a homozygous locus, the second term assumes a heterozygous locus, the third term corresponds to a homozygous locus that suffered a somatic mutation, and the last term to a heterozygous locus with a somatic mutation. See **Supplemental Material S1** for a more detailed derivation.

We then include the locus into the subset of informative positions if *P* (*M* | *c*_1_, *c*_2_, *c*_3_, *c*_4_) ≤ *K* for a suitable constant *K* (see **Supplemental Material S2** and **Supplementary Table S1**).

Filtering heterozygous loci is similar. Here, let *P* (*M*^*′*^ | *c*_1_, *c*_2_, *c*_3_, *c*_4_) be the probability that all cells have the same *heterozygous* genotype. The individual terms in **Equation 1** are identical except that the event probabilities for the multinomial distribution are 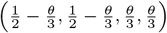. However, since heterozygous loci are three orders of magnitude fewer than homozygous loci (Bryc *et al*., 2013; Meyer *et al*., 2012) in addition to potentially being useful in haplotype phasing, we empirically determined that the following simpler and faster criteria works equally well in practice: denote the locus as informative if *c*_1_ *>* 1.5· *c*_2_, where *c*_1_ and *c*_2_ are the most frequent and the second most frequent bases at that locus, respectively (the expectation is that at a heterozygous locus *c*_1_ and *c*_2_ should not differ too much). In addition, we reject all loci for which *c*_1_ + *c*_2_ + *c*_3_ < 5.

The final set of informative loci then includes those positions that were marked as informative by both filtering steps (i.e. filtering of both homozygous and heterozygous loci). In practice, sequencing artefacts may lead to loci with unusually high coverage. For this reason, we also eliminate any loci with coverage more than two standard deviations away from the expected coverage. In addition, we also eliminate loci where *c* − *c*_1_ < 5.

### Cell-to-cell similarities

We define the similarity *s*(*i, j*) of cells *i* and *j* as the log-odds of the probability that cells *i* and *j* have the same genotype and the probability that they have different genotypes, given the corresponding sets of reads. Each of the two probabilities is then approximated as a product of probabilities of individual *overlaps* of two reads, one read from cell *i* and one read from cell *j* (**Figure 2**). Formally:

**Fig. 2.**
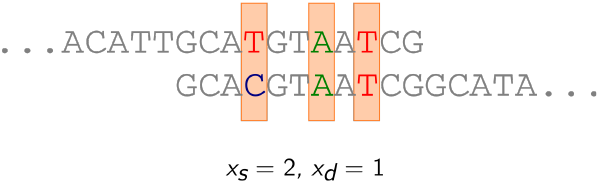
Illustration of an overlap between two reads. The orange shaded positions are the positions chosen as informative. In this example, length of the overlap is 3, the number of positions where the bases are the same, *x*_*s*_, is 2 and the number of positions where they are different, *x*_*d*_, is 1. For our purposes, an overlap is fully described by the tuple (*x*_*s*_, *x*_*d*_).

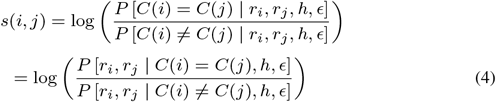

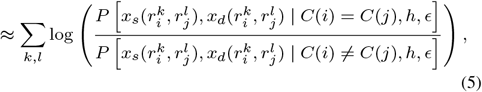

where *r*_*i*_ is the set of reads from cell 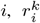 is the *k*-th read from cell *i, x*_*s*_(*p, q*) and *x*_*d*_(*p, q*) the number of matches and mismatches, respectively, between reads *p* and *q, C*(*i*) is the (true) cluster assignment of cell *i, ϵ* is the proportion of SNVs in the set of informative positions and *h* the proportion of homozygous loci in the set of informative positions (see below). In case the two cells have no overlapping reads, the similarity is by definition equal to 0 (i.e., we have no information on whether the two cells have equal or different genotypes). We assume that observing cells with the same genotype and with different genotypes has the same prior probability. (Notice that decomposing the probabilities in **Equation 4** over pairs of reads is indeed only an approximation. In particular, the decomposition in **Equation 5** would only be precise if no two reads coming from one cell were overlapping; in the opposite case, the probabilities of read pairs containing one of these overlapping reads are non-independent. However, since the per-cell coverage is so low (**Supplementary Figure S1**), the number of such non-independent pairs is negligible.)

Notice that by decomposing the probabilities over the overlaps of reads we gain information not only on the number of matches and mismatches between the two reads (i.e. information on potential differences between the two cells), but also information on haplotype phasing. Moreover, it also allows us to put more weight on longer (and hence supposedly more informative) overlaps. For example, a long overlap with only matches is an indication that the two cells might have the same genotype. A long overlap with only mismatches, on the other hand, is not a strong indication towards the cells being from different clusters – another likely scenario is that the two reads were sampled from different haplotypes and we just observe a row of heterozygous loci in different phase. As a result, overlaps with a combination of matches and mismatches are the ones most strongly suggesting the ‘different genotypes’ case (**Supplementary Figure S2**). We also show, using simulated data, that considering the number of matches and mismatches in the whole overlap of two reads provides strictly more information than considering each locus independently (**Supplementary Figure S3**).

Below we give details on the computation of **Equation 5**, under the simplifying assumptions that (1) all cells are diploid, (2) the somatic SNVs are with equal probability of type *AA*+*AB* and *AB*+*AA* (a homozygous site in cluster 1, heterozygous in cluster 2, or vice versa), and (3) the prevalence of differences between any two subclones is *μ* (see **Supplemental Material S3** for the full list of assumptions).

#### Parameters

The algorithm has three parameters: *h*, the fraction of the homozygous loci in the set of selected positions, *ϵ*, the fraction of the mutated loci in the set, and *θ*, the error rate. In our analyses, we used *h* = 0.5, *ϵ* = 0.01, and *θ* = 0.05 (the *θ* parameter has higher value than the usually reported sequencing error rate, because the set of informative positions is enriched in positions carrying sequencing errors).

*Computing the probabilities of overlaps* We define:

- *P*_*s,s*_, the probability that sequencing of two bases of the same kind results again in two bases of the same kind: 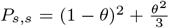 (both bases are sequenced without error, or both are misread to the same base),
- *P*_*s,d*_, the probability that sequencing of two bases of the same kind results in bases that differ from each other: *P*_*s,d*_ = 1 − *P*_*s,s*_,
- *P*_*d,s*_, the probability that two different bases are read as the same: 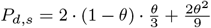 (one of the two bases is misread to the other one, or both are misread to the same base),
- *P*_*d,d*_ the probability that two different bases are sequenced as different: *P*_*d,d*_ = 1 − *P*_*d,s*_.

The probability of observing *x*_*s*_ matches and *x*_*d*_ mismatches in an overlap of length *x*_*s*_ + *x*_*d*_, assuming cells *i* and *j* have the same genotype, is now:

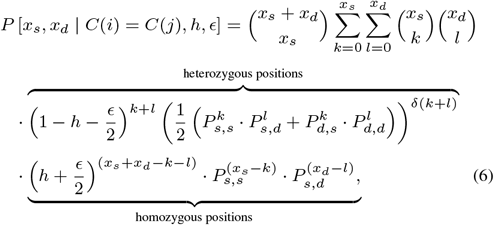

where *δ*(*x*) is a function defined as 0, if *x* = 0, and 1, otherwise. In the formula we sum over all possible combinations of (*k* + *l*) heterozygous loci and (*x*_*s*_ + *x*_*d*_ − *k* − *l*) homozygous loci; *k* of the heterozygous loci result in a match, the remaining *l* in a mismatch.

The probability of observing *x*_*s*_ matches and *x*_*d*_ mismatches assuming cells *i* and *j* are in different clusters is:

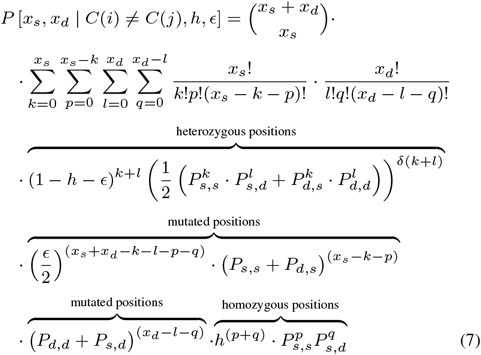

Here *k* denotes the number of heterozygous positions giving rise to a match, *l* the number of heterozygous positions giving rise to a mismatch, *p* the number of positions with the same homozygous genotype in both types of cells that give rise to a match and *q* the number of these positions that result in a mismatch.

### Clustering

We first normalize the computed similarity matrix by making sure all elements are positive: *S*^*^ = −*S* + min_*i,j*_ *s*(*i, j*). The cells are then clustered using a slight variation on spectral clustering (Ng *et al*., 2001) as follows. We compute the symmetric normalized Laplacian 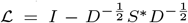 and determine its first *k* (we used *k* = 6 in all experiments in this paper) eigenvectors, corresponding to the *k* smallest eigenvalues. We then cluster into 1, 2, 3 or 4 clusters using k-means (Arthur and Vassilvitskii, 2006; Lloyd, 1982), computing the inertia values *i*_1_, *i*_2_, *i*_3_, *i*_4_ for each of the four options and the inertia gaps *g*_*k*_ = *i*_*k*_ − *i*_*k*−1_, *k* = 2, 3, 4, and define *g*_1_ := 0. The final number of clusters is max_*k*=2,3,4_ {*k* | *g*_*k*_ *>* 0.75*g*_*k*−1_}.

An important feature of spectral clustering is that it leverages the information on similarities of all pairs of cells at the same time. Thus, even in case two cells would not have any overlapping reads (the probability of which is negligibly small, see **Supplemental Material S4**), they could still be clustered based on their similarities to other cells in the data set.

One important aspect of clustering is the stopping criterion, i.e. the decision whether a specific group of cells should be divided into subclusters or not. We suggest a (to the best of our knowledge new) heuristic approach to automatically decide if the computed normalized similarity matrix *S*^*^ indicates that there are two (or more) different clusters of cells. We fit a Gaussian mixture model with 1, 2, 3 or 4 components to the smallest *k* eigenvectors of *S*^*^ and compare their likelihood using the Akaike information criterion (AIC) or the Bayesian information criterion (BIC). If the model with only one component is preferred by AIC/BIC over the models with 2, 3 or 4 components we do not split the data further. We further do not accept the split if the resulting subclone has too few cells (we used 500 in our experiments). We also require that the mean within-cluster coverage is at least 9, the lowest coverage sufficient for a reliable variant call (see **Supplemental Material S5**).

## 3 Results

### SECEDO recovers tumor subclones with average precision of 97% on simulated data

In order to test the performance of our method, we simulated a dataset consisting of 7,250 cells divided into 9 groups of various sizes: one group of healthy cells and 8 groups of tumor cells. The genome of the healthy cells was created using Varsim (Mu *et al*., 2014) based on the GRCh38 human reference genome. Common variants from dbSNP (Sherry *et al*., 2001) (3,000,000 single-nucleotide polymorphisms, 100,000 small insertions, 100,000 small deletions, 50,000 multi-nucleotide polymorphisms, 50,000 complex variants) were added to the genome. The genome of the tumor cells was built by adding 2,500 to 20,000 of both coding and non-coding SNVs (subclonal SNV fraction of 3%-27% (Dentro *et al*., 2021)), randomly chosen from the COSMIC v94 (Catalogue Of Somatic Mutations In Cancer) database (Tate *et al*., 2018), to the parent genome, in addition to 250 small insertions, 250 small deletions, 200 multi-nucleotide variants and 200 complex variants (**Figure 3**, left). Paired-end reads, with each mate of length 100 bp, were simulated using ART (Huang *et al*., 2011) at an average coverage of 0.05x per cell and with the error profile of Illumina HiSeq 2000 machines. The reads were then aligned using Bowtie 2 (Langmead and Salzberg, 2012) and filtered using Samtools (Li, 2011) to select for reads mapped only in proper pair, non-duplicate and only primary alignments.

**Fig. 3.**
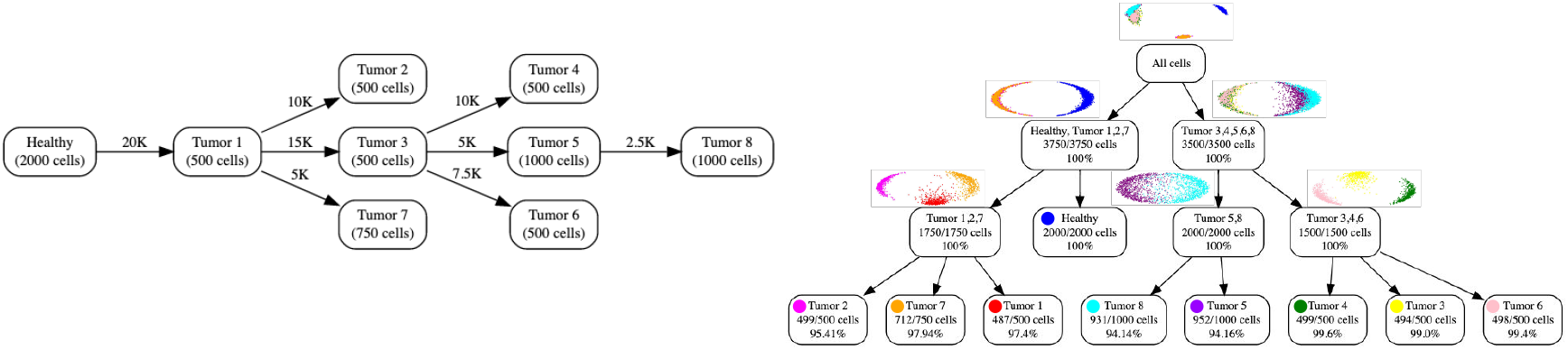
Clustering a synthetic dataset with 9 unequally sized subclones totaling 7,250 cells. Left: Theoretical phylogenetic tree of the dataset. Edge labels indicate the number of additional SNVs in each subclone relative to the parent, node labels indicate the number of cells in each subclone. Right: Recursive clustering by SECEDO. Each nodes corresponds to one SECEDO clustering step. The percentage at the bottom of each node indicates the clustering precision (correctly clustered cells relative to total cells in cluster). The scatter plots above parent nodes depict the 2nd and 3rd eigenvectors of the similarity matrix Laplacian. For leaf nodes SECEDO correctly determined that further clustering is not desirable.

For efficiency reasons, we build the pileup files used by the Bayesian filtering using our own implementation rather than existing tools that are not optimized for use on thousands of cells simultaneously (e.g. Samtools, which currently does not offer a multi-threaded pileup creation). The pileup creation, distributed on 23 commodity machines (one for each chromosome) using 20 threads each, takes about 70 minutes (down from 72 hours when using Samtools’ pileup creation on the same machines). We ran SECEDO on the resulting pileup files on an Intel(R) Xeon(R) Gold 6140 CPU @ 2.30GHz using 20 threads and 32GB of RAM. The filtering, clustering and VCF generation took 21 minutes. For the top level clustering, the filtering step kept about 1 in 16,000 loci. Somewhat counter-intuitively, the number of filtered loci approximately doubled at each level as we traveled down the clustering tree. This is due to the fact that the discriminative power of the Bayesian filtering degrades as the mean pooled coverage decreases (from 248 at the root to 20 at the leaves), such that a larger proportion of loci that are not relevant are let through. SECEDO was able to recover all 9 subclones with an average precision of 97.45% (**Figure 3**, right). Note that SECEDO is not attempting to reconstruct the evolutionary history of the tumor, but merely trying to efficiently find a grouping of cells that reflect the current subclonal structure and enable downstream tasks like variant calling. Therefore, the clustering tree reconstructed by SECEDO does not reflect the actual developmental process that gave rise to the given population of cancer cells; indeed, the SECEDO clustering tree differs from the true phylogenetic tree of the population (**Figure 3**).

In order to show the potential of the resulting clusters for somatic variant calling, we identified the most likely genotype of each cluster using a simple MAQ-based approach (Li *et al*., 2008) (**Supplemental Material S5**) and generated VCF files for each cluster against the GRCh38 human reference genome. Similarly to other variant callers that remove germline variants (Cibulskis *et al*., 2013), we then removed the ground-truth variants that were present in the healthy cells and compared the remaining SNVs against the ground truth SNVs provided by Varsim for each cluster. SECEDO was able to detect 92.11% of the somatic SNVs (vs. 77.79% when calling variants on the unclustered cells) with a 52.41% average precision (see **Supplementary Table S2**).

### SECEDO is able to correctly group cells starting at 0.03x coverage and 500 cells per cluster

One practical question of crucial importance is how to determine if, given a dataset, SECEDO will be able to correctly cluster the cells for meaningful downstream processing. To answer this question, we conducted a series of experiments to determine the conditions under which SECEDO can successfully be applied to a given dataset. There are three cluster attributes that affect SECEDO’s ability to separate cell clusters: (a) the number of cells, (b) the average per-cell coverage, and (c) the number of SNVs in which the clones differ. In order to test the interplay of these three cluster attributes, we devised a series of synthetic datasets, each consisting of 1,000 cells belonging to two groups. The sizes of the two groups were either equal (i.e. 500 cells in each group) or in ratio 1:3 (i.e. one cluster consisted of 250 cells and the other one of 750 cells). We further constructed a series of synthetic data sets consisting of 2,000 cells being split equally among two groups (i.e. 1,000 cells in each group). Then, for a given number of SNVs and given sizes of clusters, we gradually lowered the per-cell coverage until the algorithm was unable to cluster the cells correctly. The genome creation, reads simulation, and alignment were done as described in the previous section. For most parameter configurations, the currently achievable per-cell coverage of 0.05x is sufficient for SECEDO to correctly cluster the cells (see **Figure 4**). Since SECEDO is able to discriminate between balanced clusters of 1,000 cells that differ in as little as 2,500 SNVs (equivalent to an SNV prevalence of ca 8.33*·*10^−7^), the method can be applied to a wide variety of cancers, starting from those with very high mutation rates, such as melanoma (median prevalence of somatic SNVs ca 10^−5^) down to pancreatic and breast cancer (median prevalence of somatic SNVs ca 10^−6^) (Alexandrov *et al*., 2013; Lawrence *et al*., 2013). Note that there is a relationship between tumor mutational burden and SECEDO’s ability to distinguish subclones. SECEDO is able to identify complex subclonal structures (such as in **Figure ??**) in cancers with high mutational burden (e.g. melanoma), whereas in cancers with lower mutational burden (e.g. pancreatic and breast cancer) only major clones could be identified, as shown in the next section. As expected, the discriminative power of SECEDO increases with the number of cells (**Figure 4**), as well as with the per-cell coverage (**Supplementary Figure S4**), since both act as a multiplying factor for the pooled coverage.

**Fig. 4.**
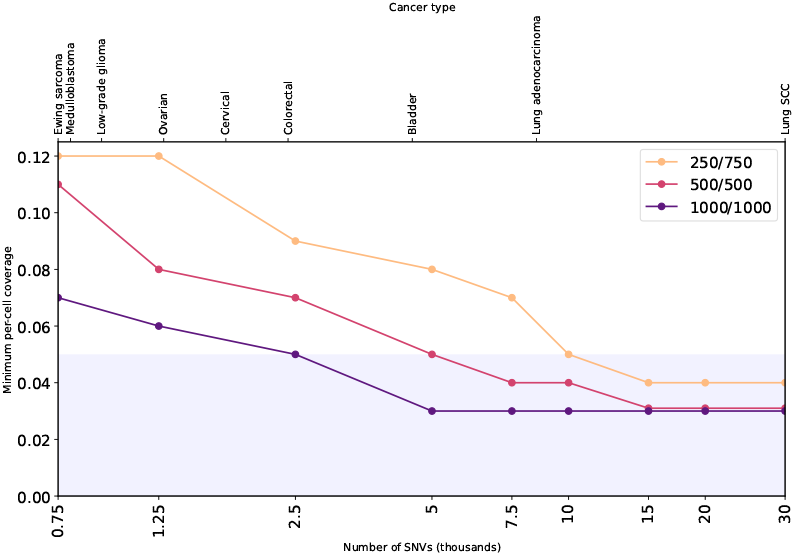
Minimum required coverage for successful clustering of sub-clones differing in the given number of SNVs, in three scenarios: clustering 1,000 cells, with a (1/4, 3/4) split, with an equal (1/2, 1/2) split, and clustering 2,000 cells with an equal split. The shaded area marks the coverage currently achievable in practice. The top labels indicate the cancer type with median mutation rate closest to the given SNV density (cancer mutation rates according to Lawrence et al. (2013)).

### SECEDO recovers dominant subclones in a breast cancer dataset, clearly outperforming state of the art

In order to test the performance of SECEDO on real data, we downloaded a publicly available 10X Genomics single-cell DNA sequencing dataset^2^ sequenced using an Illumina NovaSeq 6000 System. The dataset contains five tumor sections (labeled A to E) of a triple negative ductal carcinoma, each section containing roughly 2,000 cells (10X Genomics, 2018). The mean per-cell coverage in the data set is 0.03x, with individual coverages ranging from 0.006x to 0.086x. CHISEL, the CNV-based clustering algorithm proposed by Zaccaria and Raphael (2021), identified three dominant clones in each of the sections, except for section A, which consists mainly of healthy cells and was thus not included in our analysis.

We applied SECEDO to the four datasets corresponding to sections B,C,D, and E. The filtering step reduced the number of loci in each tumor section to roughly 1,000,000 bp (ca 0.03% of the original size); the average pooled coverage across the ≈ 2,000 cells in each dataset ranged from 45 to 55. SECEDO was able to correctly recover the three dominant clones in each of the four tumor sections. The clustering results match with high accuracy (96.68% recall, 66.59% precision) those computed by CHISEL (**Figure 5**). The scatter plots of the 2nd and 3rd eigenvector of the similarity matrix confirm that each tumor section consists of three highly separable clusters.

**Fig. 5.**
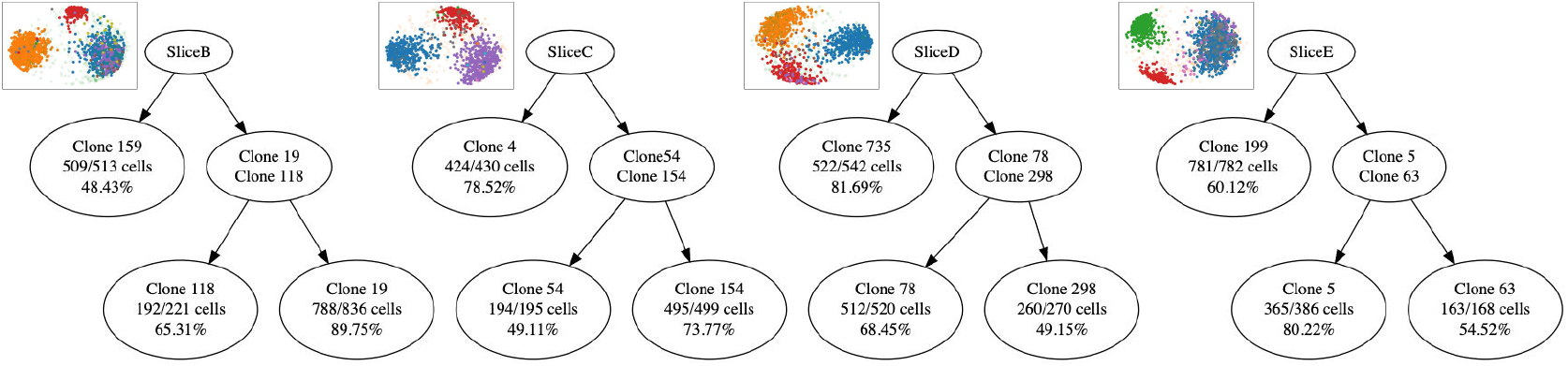
Clustering of the four tumor sections in the 10x Genomics ductal carcinoma dataset. The first row in each node denotes the cluster name; for consistency, we used the same cluster numbering as CHISEL^3^. The second row denotes the number of cells recovered by SECEDO vs the total number of cells as identified by CHISEL. The last row denotes the precision of the clustering, i.e. the percentage of cells in the SECEDO cluster that match the originally reported cluster. The lower precision values are due to the fact that cells categorized by CHISEL as “None” based on the CNV signature are assigned a category by SECEDO based on the genomic signature.

We compared SECEDO’s performance to that of SBMClone (Myers *et al*., 2020), the current state of the art in SNV-based clustering. Since SBMClone was reported to work only at coverage ≥ 0.2x, and the coverage of the breast cancer dataset is 0.03x, we created higher coverage data *in silico* by merging sequencing data from cells reported to be in the same cluster by CHISEL. In addition, SBMClone requires a matched normal sample, so we again used the clustering in CHISEL to determine the healthy cells; from the variants determined using Varscan (Koboldt *et al*., 2009), we removed all mutations that appeared in at least one healthy cell, and the remaining mutations were fed to SBMClone. SECEDO does not require a matched normal sample, so the sequencing data was used without this pre-processing. SECEDO correctly clustered (precision >96%) all cells at the original coverage (including the separation of healthy cells), and its performance remained relatively constant as coverage increased. SBMClone was able to provide an approximate clustering starting at 3-fold the original coverage, and its performance matched SECEDO’s at 7-fold the original coverage when combining data from all slices. For individual slices, SBMClone was not able to cluster the cells, irrespective of the coverage (**Figure 6**).

**Fig. 6.**
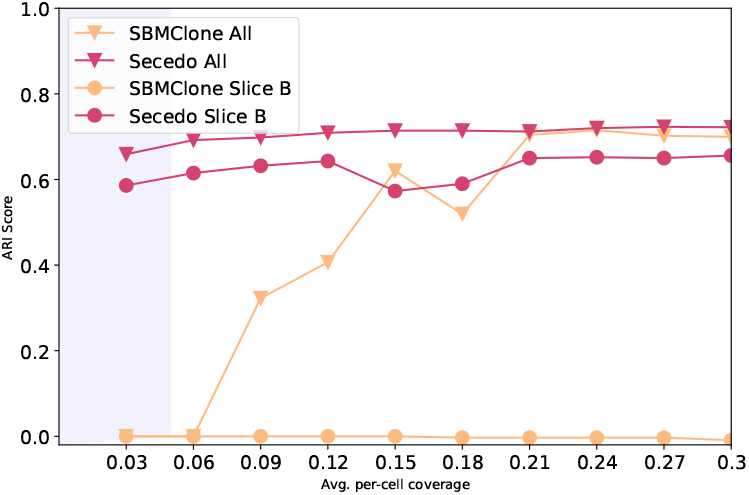
Adjusted Rand Index scores for the SECEDO and SBMClone clustering for Slice B and all slices of the breast cancer dataset at coverage ranging from 0.03x to 0.3x. Shaded area marks the average per-cell coverage achievable with current technology.

We then called SNVs on each subclone of Slice B, as identified by SECEDO, independently, and on the entire slice. In order to call SNVs, we created a Panel of Normals from the cells categorized as normal by CHISEL based on the CNV profile (Clone19 in the left-most tree of **Figure 5**). We ran MuTect 1.1.4 (Cibulskis *et al*., 2013) with the default settings, using dbSNP (Sherry *et al*., 2001) and Cosmic v94 (Tate *et al*., 2018) as priors. The number of distinct SNVs in the two tumor subclones is more than double the number of variants that were called when pooling all cells together (**Figure S5**, left). The histogram of the allelic ratio for the sublconal and global SNVs shows a significant shift to the right for the subclonal SNVs, an indication that the clustering correctly identified and separated genetically similar cells, enabling the detection of twice as many SNVs at twice the allelic ratio (**Figure S5**, right).

## 4 Discussion

We introduced SECEDO, a method that is able to correctly identify SNV-based subclones in single-cell sequencing datasets with coverage as low as 0.03x per cell. This is a significant improvement in comparison to SBMClone, the current state-of-the-art method (Myers *et al*., 2020), which, using the same data, was able to cluster the cells only after pooling data from all four data sets and artificially increasing the coverage by a factor of 7. This improvement in performance can be likely attributed to the fact that SECEDO takes into account the information on read phasing, as well as its efficient filtering of uninformative positions. We also note that unlike SBMClone, SECEDO does not require a matched normal sample for the identification of potential SNVs. We provide an efficient, well-tested, ready-to-use C++ implementation of SECEDO, which uses established data formats for both input and output, and can thus be easily incorporated into existing bioinformatics pipelines.

We demonstrated SECEDO’s applicability to currently available single-cell sequencing data and find that SECEDO correctly clustered cells on a series of synthetic and four breast cancer datasets. CNA frequencies and patterns vary significantly across cancer types (Harbers *et al*., 2021; Zack *et al*., 2013), similarly to SNV frequency. Since SECEDO does not use copy-number information to cluster cells, it can infer sub-clones even in cancer types where CNAs do not vary or where the frequency of CNAs is generally low (e.g. pancreatic neuroendocrine tumors (Dentro *et al*., 2021)). It is also notable that not all CNAs affect the SNV profile of a cell. Thus, CNA-based clustering may lead to suboptimal grouping of cells, e.g. from a variant calling perspective. SECEDO is able to group cells with similar SNV profiles irrespective of their CNA profiles. This can lead to improvements in the precision and accuracy of the variant calling. Using the clusters identified by SECEDO, we were able to recover 92.11% of the SNVs present in the synthetic data set using a simple variant caller. On Slice B of the breast cancer data set, the number and the confidence of the called SNVs more than doubled after clustering using SECEDO, compared to calling variants on the entire slice.

While SECEDO enables accurate cell-clustering and variant calling, there are a number of areas for future improvement. First, SECEDO currently only uses single-nucleotide substitutions to cluster cells, which are known to be the most common type of mutations in adult and childhood cancers (Gröbner *et al*., 2018; Lawrence *et al*., 2014; Ma *et al*., 2018). We expect that the clustering accuracy could be further improved if e.g. short insertions and deletions were additionally used. Second, the smallest subclones that SECEDO was able to detect had ≈200 cells. However, as technology inevitably improves and the sequencing coverage increases, SECEDO’s resolution and variant calling quality will also proportionally increase.

We hope that SECEDO will facilitate new types of analyses and form the basis for future methodological development in the field of cancer research and treatment outcome prognosis.

## Acknowledgements

We would like to thank Ximena Bonilla for her constructive feedback on the manuscript. This work was supported by ETH core funding to G.R. (funding D.D., S.S., A.K., KV.L.).

## Footnotes

1 https://www.10xgenomics.com/resources/datasets/

2 https://www.10xgenomics.com/resources/datasets/

3 Available at https://github.com/raphael-group/chisel-data/

