## Supplementary Material for "SECEDO: SNV-based subclone detection using ultra-low coverage single-cell DNA sequencing"

#### S1 Approximating the probability of the evidence

In this section we show how we approximate the probability of the evidence $P(c_1, c_2, c_3, c_4)$, where $c_i$ represents the number of bases piled up at a certain locus, sorted by most frequent to least frequent, $\mathbf{C} = (c_1, c_2, c_3, c_4)$, $c = c_1 + c_2 + c_3 + c_4$ is the pooled coverage at the locus, and by $C_1, \ldots, C_4$ we denote the corresponding bases. Similarly, denote by $a_1, \ldots, a_4$ the (unknown) counts of $C_1, \ldots, C_4$ among the true alleles, from which the observations were made. For example, the true sequenced alleles might be AAACAC, so $a_1 = 4$, $a_2 = 2$, $a_3 = a_4 = 0$, but due to a sequencing error the observed bases are AAGCAC, so $c_1 = 3$, $c_2 = 2, c_3 = 1$, $c_4 = 0$ and $C_1 = A$, $C_2 = C$, $C_3 = G$, $C_4 = T$.

The goal is to approximate $P(\mathbf{C})$. This is equal to the sum over all possible (unknown) combinations of the true alleles:

$$P(\mathbf{C}) = \sum_{\mathbf{A}} P(\mathbf{C} \mid \mathbf{A}) P(\mathbf{A}),$$

where $\mathbf{A} = (a_1, a_2, a_3, a_4)$. The first term quantifies the probability that sequencing $a_1$ bases $C_1$, $a_2$ bases $C_2$, etc. will result in the observation of $c_1$ bases $C_1$, $c_2$ bases $C_2$, etc., the second is the prior on the allele combination $\mathbf{A}$.

The sum above can be approximated by summing over the following four combinations of alleles:

1. All true alleles are $C_1$ (i.e. $a_1 = c$, $a_2 = a_3 = a_4 = 0$), i.e. all cells have the same homozygous genotype. This has prior probability $p_{hom}$ and $P(\mathbf{C} \mid \mathbf{A})$ is equal to multinomial probability of $(c_1, c_2, c_3, c_4)$ with probabilities $(1 - \theta, \frac{\theta}{3}, \frac{\theta}{3}, \frac{\theta}{3})$, so we get:

$$P(\mathbf{C} \mid \mathbf{A}_{hom}) P(\mathbf{A}_{hom}) = p_{hom} \frac{c!}{c_1! c_2! c_3! c_4!} (1 - \theta)^{c_1} \left(\frac{\theta}{3}\right)^{c_2 + c_3 + c_4}$$

2. The locus is heterozygous $C_1 C_2$, which has prior $p_{het}$. The most likely case is that in reality there were $c_1$ occurrences of $C_1$, $c_2$ of $C_2$ and then some combination of $c_3 + c_4$ of either $C_1$ or $C_2$ (the exact combination is unknown, but also irrelevant for the computation). We can then compute $P(\mathbf{C} \mid \mathbf{A}_{het}) P(\mathbf{A}_{het})$ as

$$P(\mathbf{C} \mid \mathbf{A}_{het}) P(\mathbf{A}_{het}) = p_{het} \frac{c!}{c_1! c_2! c_3! c_4!} \left(\frac{1}{2} - \frac{\theta}{3}\right)^{c_1 + c_2} \left(\frac{\theta}{3}\right)^{c_3 + c_4}$$

3. The locus was originally homozygous $C_1C_1$ but one haplotype suffered a somatic mutation, so the new genotype is $C_1C_2$ in some cells. This has prior $p_{hom}p_{mut}$. Assuming the proportion of tumor cells is 0.5, the most likely case is that in reality there were $c_1$ occurrences of $C_1$, $c_2$ occurrences of $C_2$ (about 3 times less than $c_1$) and then some combination $c_3 + c_4$ of either $C_1$ or $C_2$. Again, we don't know how many of the sequence errors were originally $C_1$ and how many $C_2$, and it doesn't matter. The multinomial probabilities are in this case equal to $(\frac{3}{4} - \frac{2}{3}\theta, \frac{1}{4}, \frac{\theta}{3}, \frac{\theta}{3})$ and $P(\mathbf{C} \mid \mathbf{A}_{hom,mut})P(\mathbf{A}_{hom,mut})$ can be computed as

$$P(\mathbf{C} \mid \mathbf{A}_{hom,mut})P(\mathbf{A}_{hom,mut}) = p_{hom}p_{mut}\frac{c!}{c_1!c_2!c_3!c_4!}\left(\frac{3}{4} - \frac{2\theta}{3}\right)^{c_1}\left(\frac{1}{4}\right)^{c_2}\left(\frac{\theta}{3}\right)^{c_3+c_4}$$

4. The locus was heterozygous and one somatic mutation happened, i.e. some cells have genotype $C_1C_2$ and some have genotype $C_1C_3$. This has prior $p_{het}p_{mut}$. Assuming again equally sized subclones, we have:

$$P(\mathbf{C} \mid \mathbf{A}_{\mathrm{mut}})P(\mathbf{A}_{\mathrm{mut}}) = p_{het}p_{mut}\frac{c!}{c_1!c_2!c_3!c_4!}\left(\frac{1}{2} - \frac{\theta}{3}\right)^{c_1}\left(\frac{1}{4}\right)^{c_2+c_3}\left(\frac{\theta}{3}\right)^{c_4}$$

We neglect all the other possibilities (e.g., the possibility of the locus undergoing more than one somatic mutation), as they have very low prior probability.

To get $P(c_1, c_2, c_3, c_4)$, we then simply sum over all four possibilities:

$$\begin{aligned}P(c_1, c_2, c_3, c_4) \approx \frac{c!}{c_1!c_2!c_3!c_4!}\Bigg[&p_{hom}(1-\theta)^{c_1}\left(\frac{\theta}{3}\right)^{c_2+c_3+c_4} + \\ &+ p_{het}\left(\frac{1}{2} - \frac{\theta}{3}\right)^{c_1+c_2}\left(\frac{\theta}{3}\right)^{c_3+c_4} + \\ &+ p_{hom}p_{mut}\left(\frac{3}{4} - \frac{2\theta}{3}\right)^{c_1}\left(\frac{1}{4}\right)^{c_2}\left(\frac{\theta}{3}\right)^{c_3+c_4} + \\ &+ p_{het}p_{mut}\frac{c!}{c_1!c_2!c_3!c_4!}\left(\frac{1}{2} - \frac{\theta}{3}\right)^{c_1}\left(\frac{1}{4}\right)^{c_2+c_3}\left(\frac{\theta}{3}\right)^{c_4}\Bigg]\end{aligned}$$

#### S2 Finding the optimal values of K for filtering uninformative loci

To find the optimal value of the constant $K$, we simulated a data set as follows: We assumed that there are only two types of cells in the sample ('healthy' and 'tumor'), that sequencing errors are unbiased and happen independently of each other with rate $\theta = 0.001$, that the somatic SNVs can only be of type $AA$+$AB$ (i.e., a heterozygous mutation in tumor cells), $AB$+$AA$ (i.e., loss of heterozygosity in tumor cells) or $AA$+$BB$ (i.e., a homozygous mutation) and that these three types of somatic mutations are equally likely. We also assumed that the coverage values at individual sites follow a Poisson distribution with a specified mean and that all loci are diploid. For different sample compositions (proportion of tumor cells in the sample ranging from 10 to 50%; notice that by symmetry the case with e.g. 60% tumor and 40% healthy cells is equivalent to 40% tumor and 60% healthy), we simulated 100,000 results for each of the following scenarios:

- all cells have the same homozygous genotype – the counts follow a multinomial distribution with probabilities $\left(1-\theta,\ \frac{\theta}{3},\ \frac{\theta}{3},\ \frac{\theta}{3}\right)$
- all cells have the same heterozygous genotype – the counts follow a multinomial distribution with probabilities $\left(\frac{1}{2}-\frac{\theta}{3},\ \frac{1}{2}-\frac{\theta}{3},\ \frac{\theta}{3},\ \frac{\theta}{3}\right)$
- a somatic mutation, healthy cells have genotype $AA$, tumor cells $AB$ – the counts follow a multinomial distribution with probabilities $\left((f_h+\frac{1}{2}f_t)(1-\theta)+\frac{1}{2}\frac{\theta}{3}f_t,\ \frac{1}{2}f_t(1-\theta)+(f_h+\frac{1}{2}f_t)\frac{\theta}{3},\ \frac{\theta}{3},\ \frac{\theta}{3}\right)$
- a somatic mutation, healthy cells have genotype $AB$, tumor cells $AA$ – the counts follow a multinomial distribution with probabilities $\left((f_t+\frac{1}{2}f_h)(1-\theta)+\frac{1}{2}\frac{\theta}{3}f_h,\ \frac{1}{2}f_h(1-\theta)+(f_t+\frac{1}{2}f_h)\frac{\theta}{3},\ \frac{\theta}{3},\ \frac{\theta}{3}\right)$
- a somatic mutation, healthy cells have genotype $AA$, tumor cells $BB$ – the counts follow a multinomial distribution with probabilities $\left(f_h(1-\theta)+f_t\frac{\theta}{3},\ f_t(1-\theta)+f_h\frac{\theta}{3},\ \frac{\theta}{3},\ \frac{\theta}{3}\right)$,

where $f_h$ is the proportion of healthy cells in the sample and $f_t = 1 - f_h$ the proportion of tumor cells.

The optimal value of $K$ was then defined as the value maximizing Youden's J [3]:

$$J = \text{sensitivity} + \text{specificity} - 1 = \frac{TP}{TP+FN} + \frac{TN}{TN+FP} - 1.$$

The optimal values of $K$ for different combinations of pooled coverage and sample composition (proportion of healthy and tumor cells in the sample) are given in **Supplementary Table S1**.

#### S3 Assumptions for computing the cell-to-cell similarities

We make the following simplifying assumptions:

1. The prevalence of differences between any two subclones is $\mu$.

2. All cells are diploid.

3. The somatic SNVs are with equal probability of type $AA$+$AB$ (a homozygous site in cluster 1, heterozygous in cluster 2) and $AB$+$AA$ (a heterozygous site in cluster 1, homozygous in cluster 2); in particular, we do not take into account homozygous SNVs ($AA$+$BB$) or mutations that involve more than two different alleles (e.g. $AB$+$AC$) [1].

4. The mutated and germline heterozygous loci are distributed randomly and independently in the genome; in particular, they do not tend to cluster together.

5. Sequencing errors happen independently at each sequenced base with probability $\theta$ and are unbiased (i.e. all types of substitutions are equally likely).

6. We run the clustering on the set of informative positions; in the set there are loci of three types:

   - loci that carry a somatic SNV,
   - loci that have the same homozygous genotype in all cells, and were not discarded during the previous step only because of sequencing errors, and
   - loci that have the same heterozygous genotype in all cells, and were not discarded during the previous step e.g. because of preferential amplification.

   We denote the frequency of the homozygous loci in the data set by $h$ and the frequency of the mutated loci by $\epsilon$; the frequency of the germline heterozygous loci is then $(1 - h - \epsilon)$.

#### S4 Probability that two cells do not have any informative read overlaps

Let $r$ be the average proportion of loci covered by at least one read in a single cell (for very low values of mean per-cell coverage, $r$ is roughly equal to the coverage) and $I$ the number of informative bases as identified by the Bayesian filtering step. The probability that a single informative locus is not covered by a read in both of two given cells is then

$$\left(1 - r^2\right)$$

and, approximating the informative loci as independent of each other, the probability that none of the informative bases is covered by a read in both cells is roughly equal to

$$\left(1 - r^2\right)^I .$$

For $r = 0.05$ and $I = 2 \cdot 10^6$ (i.e., the frequency of informative loci in the genome is roughy 1 in 16,000, as observed in our experiments), the probability is in the order of $10^{-2174}$.

### S5 Variant calling: the MAQ approach

In this section we describe a simple variant calling approach inspired by the MAQ algorithm [2]. We start by creating a pseudo-bulk sample by pooling together data from all cells in one cluster. Then we call the most likely genotype for each cluster separately.

More precisely, for each site $s$ with sufficient coverage (see below), we calculate the posterior distribution of all possible genotypes and call the most likely genotype:

$$G_j[s] = \max_{g \in \mathcal{G}} P\left(g \mid x_j^A[s], x_j^C[s], x_j^G[s], x_j^T[s], \theta\right), \tag{S1}$$

where by $\mathcal{G}$ we denote the set of all possible genotypes (AA, AC, AG, AT, CC, etc.), by $x_j^A[s], x_j^C[s], x_j^G[s]$ and $x_j^T[s]$ the number of A's, C's, G's and T's, resp., read at position $s$ in cluster $j$, and by $\theta$ we denote the error rate. Using Bayes theorem we can rewrite **Equation S1** as

$$\begin{aligned} G_j[s] &= \max_{g \in \mathcal{G}} \frac{P\left(x_j^A[s], x_j^C[s], x_j^G[s], x_j^T[s] \mid g, \theta\right) P(g)}{P\left(x_j^A[s], x_j^C[s], x_j^G[s], x_j^T[s] \mid \theta\right)} \\ &= \max_{g \in \mathcal{G}} P\left(x_j^A[s], x_j^C[s], x_j^G[s], x_j^T[s] \mid g, \theta\right) P(g). \end{aligned} \tag{S2}$$

The probability of $x_j^A[s], x_j^C[s], x_j^G[s], x_j^T[s]$ given genotype $g$ and error rate $\theta$ follows a multinomial distribution with probabilities $\left(1-\theta, \frac{\theta}{3}, \frac{\theta}{3}, \frac{\theta}{3}\right)$ if $g$ is homozygous and $\left(\frac{1}{2}-\frac{\theta}{3}, \frac{1}{2}-\frac{\theta}{3}, \frac{\theta}{3}, \frac{\theta}{3}\right)$ if $g$ is heterozygous. The priors are 1 if $g$ is homozygous and 0.001 if $g$ is heterozygous, as chosen by the authors of MAQ [2].

Notice that the genotype maximizing **Equation S1** does not need to be unique. If such a situation happens, we do not call any genotype for the given site.

Variant calling is then simply done by comparing the genotypes called in individual clusters.

To decrease the number of false calls, we further only call genotypes for sites with coverage at least 9 (in the given cluster) because at low-coverage sites there is a high chance of an incorrect genotype call. In particular, if the coverage of a site is smaller or equal to 3 and assuming $\theta = 0.001$, it is actually impossible to call a heterozygous genotype using the algorithm described above. Similarly, if the site is covered by 4 reads, the probability of correctly calling a heterozygous genotype is only 37.4%. Hence, we decided to call genotypes only for sites where the coverage is large enough, such that the theoretical chance of correctly calling both homozygous and heterozygous genotype is at least 95%.

We denote coverage by $c$ (meant is the within-cluster pooled coverage) and sequencing error rate by $\theta$, $\theta = 0.001$. We assume all cells are assigned to the correct cluster, i.e. all cells within one cluster have the same genotype. We want to find minimum $c$ such that it holds both

$$P(\text{correct genotype call} \mid c, \text{true genotype homozygous}) \geq 0.95$$

and

$$P(\text{correct genotype call} \mid c, \text{true genotype heterozygous}) \geq 0.95.$$

(By 'correct genotype call' we mean here and everywhere below 'correct and unique'.) Notice that

$$\begin{aligned} &P(\text{correct genotype call} \mid c, \text{true genotype homozygous}) \\ &\quad = \sum_{B \in \{A,C,G,T\}} P(\text{correct genotype call} \mid c, \text{true genotype is } BB) \cdot P(BB \mid \text{true genotype homozygous}) \\ &\quad = P(\text{correct genotype call} \mid c, \text{true genotype is } XX), \end{aligned} \tag{S3}$$

for any $X \in \{\mathrm{A, C, G, T}\}$, since we assume uniform prior on all homozygous genotypes and non-biasness of sequencing errors. Similarly,

$$P(\text{correct genotype call} \mid c, \text{true genotype heterozygous}) = P(\text{correct genotype call} \mid c, \text{true genotype is } XY) \tag{S4}$$

for any $X, Y \in \{\text{A,C,G,T}\}, X \neq Y$.

**Equation S3** can be rewritten as a sum over all possible results $\mathcal{D}$, where a 'result' is an ordered quadruple $(x_A, x_C, x_G, x_T)$, giving the number of sequenced A's, C's, G's and T's at the chosen locus:

$$P(\text{correct genotype call} \mid c, \text{true genotype is } XX) = \sum_{d \in \mathcal{D}} P(d \mid c, \text{true genotype is } XX) \mathbb{I}\left(\text{correct genotype call} \mid d\right), \tag{S5}$$

where by $\mathbb{I}(\cdot)$ we denote the indicator function. Similarly

$$P(\text{correct genotype call} \mid c, \text{true genotype is } XY) = \sum_{d \in \mathcal{D}} P(d \mid c, \text{true genotype is } XY)\, \mathbb{I}\,(\text{correct genotype call} \mid d)\,. \tag{S6}$$

The probability of a particular result $d$ given the true genotypes follows a multinomial distribution with probabilities $p = \left(1-\theta, \frac{\theta}{3}, \frac{\theta}{3}, \frac{\theta}{3}\right)$ (up to ordering) in case of a homozygous genotype, and $p = \left(\frac{1}{2}-\frac{\theta}{3}, \frac{1}{2}-\frac{\theta}{3}, \frac{\theta}{3}, \frac{\theta}{3}\right)$ in case of a heterozygous genotype (again up to ordering).

We numerically computed the probabilities given by **Equation S5** and **Equation S6** for coverage ranging from 1 to 20, always enumerating all possible results. The resulting probabilities for homozygous and heterozygous genotypes are plotted in **Supplementary Figure S6**. The smallest $c$ such that both the probabilities are at least 95%, is $c = 9$.

### Supplementary tables

Table S1: Optimal values for pre-processing threshold $K$, for different combinations of mean pooled coverage and sample composition. The simulation was performed as described in **Supplemental Material S2**, the coverage was sampled independently for each site from a Poisson distribution with the specified mean.

| | Proportion of tumour cells | | | | | | | | |
|---|---|---|---|---|---|---|---|---|---|
| Mean pooled coverage | 0.1 | 0.2 | 0.3 | 0.4 | 0.5 | 0.6 | 0.7 | 0.8 | 0.9 |
| 10 | -1.729 | -1.709 | -1.561 | -1.561 | -1.601 | -1.561 | -1.709 | -1.561 | -1.729 |
| 20 | -1.390 | -1.388 | -1.390 | -1.394 | -1.394 | -1.394 | -1.389 | -1.389 | -1.390 |
| 30 | -1.386 | -1.386 | -1.386 | -1.386 | -1.386 | -1.386 | -1.386 | -1.386 | -1.386 |
| 40 | -1.386 | -1.386 | -1.386 | -1.386 | -1.386 | -1.386 | -1.386 | -1.386 | -1.386 |
| 50 | -1.386 | -1.386 | -1.386 | -1.386 | -1.386 | -1.386 | -1.386 | -1.386 | -1.386 |
| 60 | -1.386 | -1.386 | -1.386 | -1.386 | -1.399 | -1.386 | -1.386 | -1.386 | -1.386 |
| 70 | -1.386 | -1.386 | -1.386 | -1.388 | -3.673 | -1.389 | -1.386 | -1.386 | -1.386 |
| 80 | -1.386 | -1.386 | -1.386 | -1.388 | -11.48 | -1.389 | -1.388 | -1.386 | -1.386 |
| 90 | -1.386 | -1.386 | -1.388 | -11.482 | -27.492 | -11.481 | -1.388 | -1.386 | -1.386 |
| 100 | -1.386 | -1.386 | -1.388 | -27.492 | -43.502 | -3.592 | -1.388 | -1.386 | -1.386 |
| 110 | -1.386 | -1.388 | -3.592 | -19.486 | -43.502 | -27.492 | -1.388 | -1.388 | -1.386 |
| 120 | -1.386 | -1.388 | -11.481 | -27.492 | -75.524 | -35.497 | -11.481 | -1.388 | -1.386 |
| 130 | -1.386 | -1.388 | -19.486 | -51.508 | -59.513 | -43.502 | -3.592 | -1.388 | -1.386 |
| 140 | -1.386 | -1.388 | -11.481 | -59.513 | -83.529 | -51.508 | -11.481 | -1.388 | -1.386 |
| 150 | -1.386 | -1.388 | -35.497 | -75.524 | -91.535 | -51.508 | -27.492 | -1.388 | -1.386 |
| 160 | -1.386 | -3.592 | -35.497 | -67.519 | -123.556 | -67.519 | -43.502 | -3.592 | -1.386 |
| 170 | -1.386 | -3.592 | -35.497 | -91.535 | -131.562 | -99.540 | -43.502 | -3.592 | -1.388 |
| 180 | -1.388 | -11.481 | -59.513 | -115.551 | -139.56 | -99.540 | -35.497 | -11.481 | -1.388 |
| 190 | -1.388 | -11.481 | -67.519 | -123.55 | -163.583 | -107.54 | -59.513 | -19.486 | -1.388 |
| 200 | -1.388 | -19.486 | -51.508 | -107.545 | -163.583 | -107.545 | -75.524 | -3.592 | -1.388 |

Table S2: Precision and recall for the variants detected by SECEDO in each subclone of the synthetic dataset in **Figure 3**.

| Subclone | Recall | Precision |
|---|---|---|
| Tumor1 | 92.86% | 36.21% |
| Tumor2 | 91.48% | 43.10% |
| Tumor3 | 92.09% | 46.52% |
| Tumor4 | 91.58% | 53.11% |
| Tumor5 | 92.93% | 67.46% |
| Tumor6 | 92.25% | 51.70% |
| Tumor7 | 92.05% | 53.02% |
| Tumor8 | 91.60% | 68.17% |

### Supplementary figures

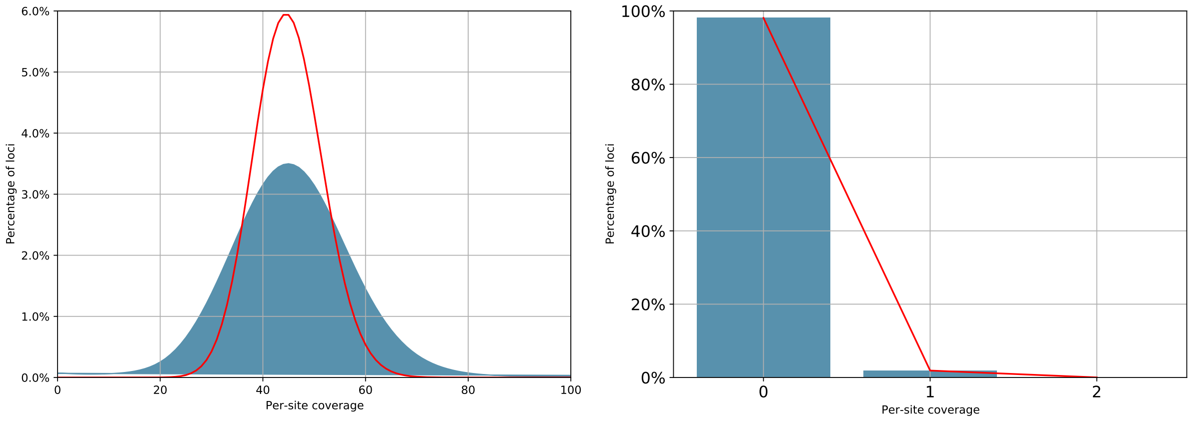


Figure S1: Coverage for Slice B of the breast cancer dataset. Left: Histogram of the per-site coverage pooled over all cells (in blue) and the theoretical Poisson distribution with the same mean (red line) Right: Histogram of the per-site coverage for individual cells. The red line is the theoretical Poisson fit.

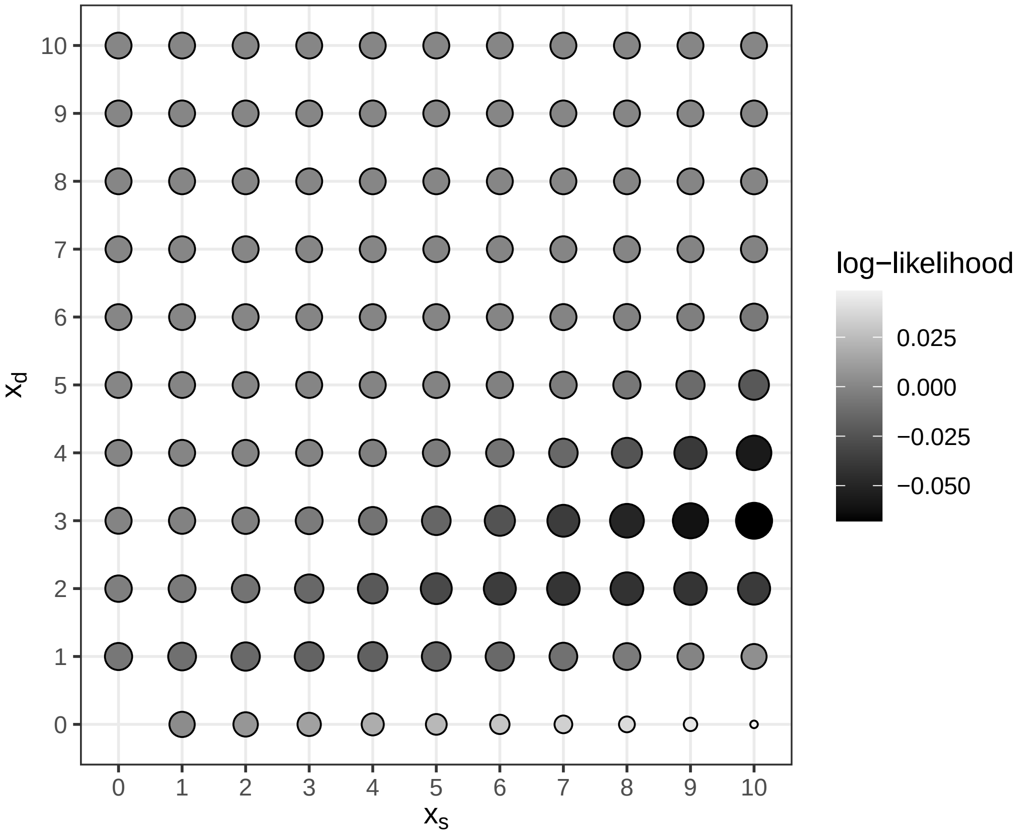


Figure S2: Informativeness of different combinations of number of matches ($x_s$) and mismatches ($x_d$) in a read overlap, computed as $\frac{P[x_s,x_d|C(i)=C(j),h,\epsilon]}{P[x_s,x_d|C(i)\neq C(j),h,\epsilon]}$. Darker and larger points denote more informative combinations. Computed using $\theta = 0.05$, $\epsilon = 0.01$, $h = 0.5$ (the same values of parameters as used for the analyses of the simulated and real data sets in the manuscript).

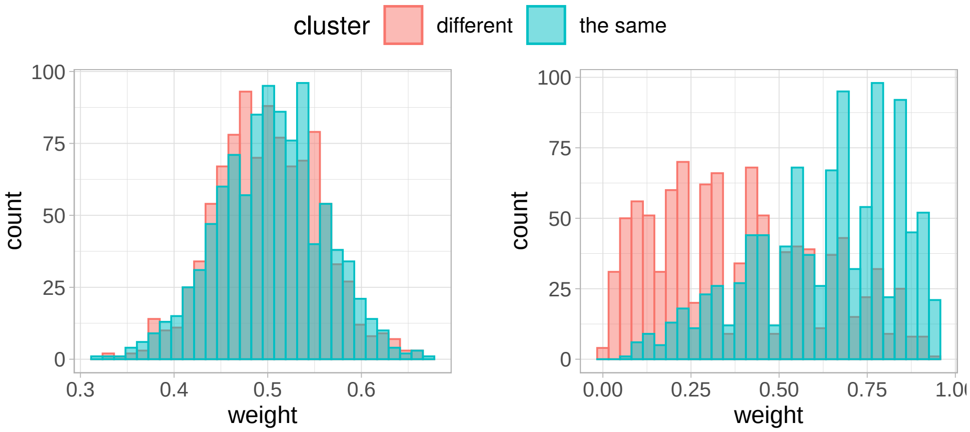


(a) Each locus treated independently (b) Overlap structure taken into account

Figure S3: Considering the overlaps as a whole provides more information than considering each locus independently. In each simulation, we simulated 550 read overlaps of length 2, assuming $\epsilon = 10^{-2}$, $h = 0$, $\theta = 10^{-3}$. The resulting weight, computed as $\frac{P[x_s,x_d|C(i)=C(j),h,\epsilon]}{P[x_s,x_d|C(i)\neq C(j),h,\epsilon]}$, was computed treating each locus independently (a) or considering the whole overlaps (b). In both cases there were 1000 simulations. The simulated data used in (a) and (b) were the same.

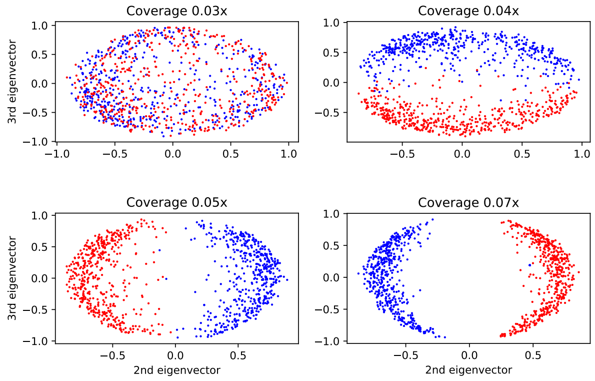


Figure S4: Scatter plots of the 2nd and 3rd eigenvectors of the similarity matrix Laplacian for increasing coverage values (synthetic data with 1,000 cells, 500 in each group; 5,000 SNVs). The higher the coverage, the clearer the separation between the two clusters.

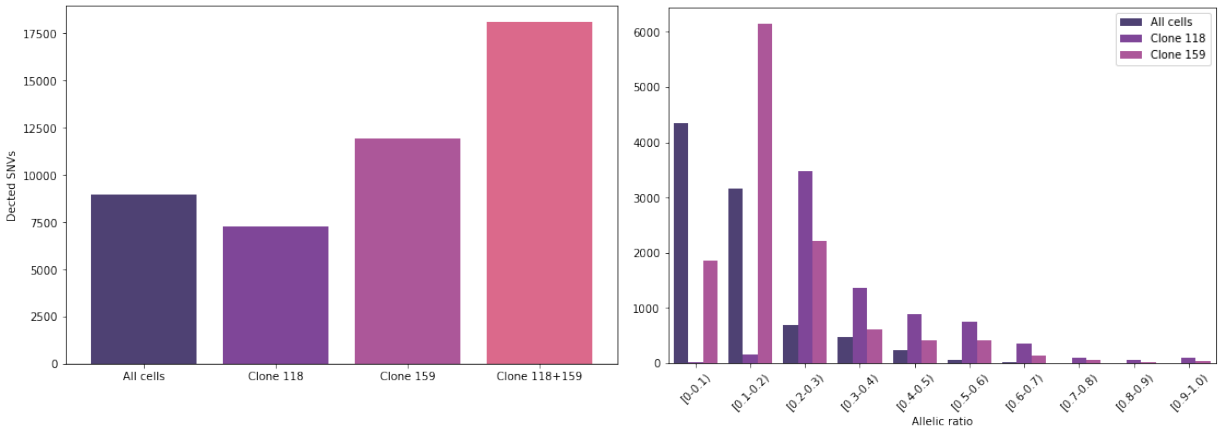


Figure S5: **Left:** SNVs detected on Slice B of the breast cancer dataset by running Mutect with the default settings. The middle columns show the number of SNVs reported when running Mutect on each of the two cancer subclones separately, as detected by SECEDO. The last column shows the number of SNVs reported by Mutect when merging the two cancer subclones, while the significantly lower value in the first column represents the number of SNVs reported by Mutect when run against all cells in the tumor slice (including the healthy cells). **Right:** Histogram of the allelic ratio for the SNVs detected by applying MuTect to the two tumor subclones and to all cells in slice B of the breast cancer dataset. The shift to the right in the allelic ratio indicates that the clustering increased the tumor purity.

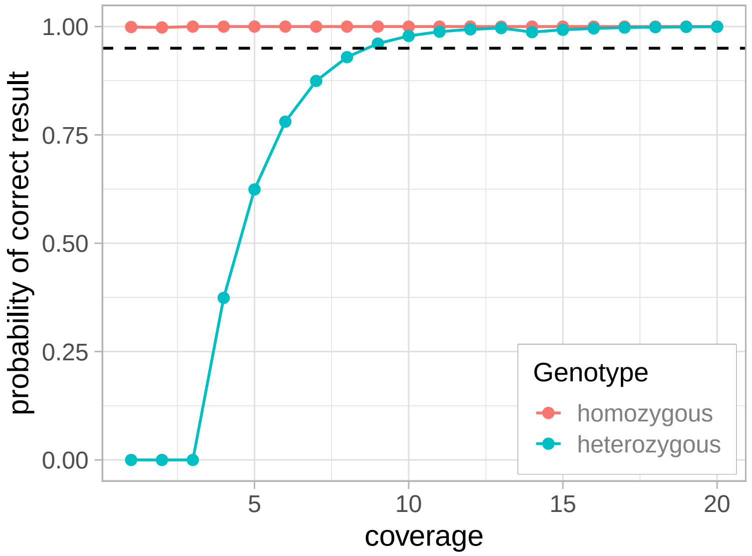


Figure S6: Probability of calling a correct and unique genotype for different values of the (within-cluster) coverage, assuming the true genotype is homozygous or heterozygous. The dashed line denotes probability 95%. See **Supplemental Material S5** for details.
